# The lipid landscape shapes the immunomodulatory potential of fluoxetine in macrophages

**DOI:** 10.64898/2026.08.25.746975

**Authors:** Jill Grondelaers, Ashleigh Jimenez-Lemus, Lieve Temmerman, Erik AL Biessen, Ronit Shiri-Sverdlov, Emiel van der Vorst, Tom Houben

## Abstract

Treatment-resistant depression (TRD) affects approximately one-third of depressed patients, yet the molecular mechanisms underlying this therapeutic non-responsiveness remain unclear. Pharmacological antidepressants, such as the selective serotonin reuptake inhibitor (SSRI) fluoxetine, exert immunomodulatory effects, partially by shifting macrophages towards an anti-inflammatory phenotype. Clinical aberrations in lipid metabolism have been associated with fluoxetine non-responsiveness in depressed populations. As macrophage polarization is highly sensitive to changes in lipid metabolism, pathological alterations in lipid metabolism may directly interfere with the therapeutic efficacy of SSRIs such as fluoxetine. However, how metabolic and immunomodulatory effects of antidepressants relate to each other in the context of TRD remains largely unexplored. We studied the interplay between fluoxetine’s immunomodulatory capacity and the macrophage lipid landscape. Human monocyte-derived macrophages (MoDMs) and murine bone marrow-derived macrophages (BMDMs) were utilized as experimental models to evaluate these localized immunometabolic effects. Under baseline conditions in wild-type macrophages, the characteristic anti-inflammatory effect of fluoxetine coincided with distinct intracellular lipid accumulation. Conversely, disrupting this lipid environment yielded opposite immunological outcomes. BMDMs deficient in the low- density lipoprotein receptor (*Ldlr^-/-^*) or wild-type BMDMs exposed to inflammatory oxidized phosphocholine-containing phospholipids (OxPLs) failed to undergo anti-inflammatory polarization and exhibited a robust pro-inflammatory response upon fluoxetine treatment instead. Collectively, these data demonstrate a critical link between the macrophage lipid landscape and fluoxetine’s immunomodulatory efficacy. These findings suggest that deficiencies in the endogenous LDLR pathway and exposure to circulating lipid peroxidation products can modulate the immunological response to fluoxetine. Our observations highlights microenvironmental lipid stress as a potential contributor to the underlying biology of antidepressant resistance in TRD.

## INTRODUCTION

Depression is a globally prevalent mood disorder that carries a significant socio-economic burden and is expected to become the world’s leading cause of disease-related disability by 2030 [1, 2]. While clinical management of severe depression is increasingly multimodal, frontline treatment remains centered on pharmacological therapies such as selective serotonin reuptake inhibitors (SSRIs) [3]. Yet, approximately one-third of patients fail to achieve remission following multiple therapeutic trials with pharmacological antidepressants [4], a phenomenon referred to as treatment-resistant depression (TRD) [5]. Therefore, identifying the mechanisms underlying treatment resistance is fundamental to overcoming current barriers to therapeutic success.

Aligned with the inflammation hypothesis of depression, previous research has suggested that pharmacological antidepressants exert part of their therapeutic effect by promoting the polarization of macrophages into a quiescent, anti-inflammatory state [6–8]. In this regard, macrophage plasticity is critically shaped by its lipid landscape, in which the balance of fatty acids, the composition of membrane phospholipids, and the intracellular handling of cholesterol collectively modulate inflammatory signaling pathways, thereby acting as metabolic determinants of the macrophage’s polarization state [9].

Parallel to these cellular dynamics, previous findings have shown a link between the use of pharmacological antidepressants and cardiometabolic parameters [10, 11]. One observation demonstrated that elevated plasma cholesterol levels, a hallmark of low-density lipoprotein receptor (LDLR) deficiency, were a significant predictor of non-responsiveness to the SSRI fluoxetine [12], a finding later confirmed for the antidepressant nortriptyline [13]. Relevantly, the LDLR serves as a critical endogenous metabolic gatekeeper, where its role in maintaining macrophage cholesterol homeostasis [14] is fundamentally linked to the control of the cell’s immune state [15, 16]. Collectively, these observations suggest that the functional integrity of the LDLR pathway could be involved with the immunomodulatory success of antidepressant treatment.

Strikingly, elevations in plasma cholesterol levels are also a hallmark of obesity [17], cardiovascular disease [18], and metabolic syndrome [19], disorders that are frequently comorbid with TRD. Notably, hypercholesterolemia challenges the functional integrity of lipid metabolism by exposing cells to pro-oxidant stressors. Prominent among these pro- oxidant stressors are lipotoxic mediators derived from lipid peroxidation, such as oxidized phosphocholine (PC)-containing phospholipids (OxPLs) [20]. OxPLs are known to engage and disrupt inflammatory signaling pathways in cardiometabolic disorders [21, 22]. Given that macrophage polarization is highly sensitive to its lipid environment, OxPLs represent a relevant component that could fundamentally undermine the immunomodulatory mechanisms of pharmacological antidepressants.

Hence, while pharmacological antidepressants exert metabolic and immunomodulatory effects, how these processes relate to each other in the context of TRD remains largely unexplored. In this study, we investigated the interplay between fluoxetine’s immunomodulatory effect and the lipid landscape of macrophages. We first characterized the metabolic remodeling induced by fluoxetine in wild-type human monocyte-derived macrophages (MoDMs) and murine bone marrow–derived macrophages (BMDMs). Subsequently, we examined the influence of the LDLR and also tested the impact of inflammatory OxPLs on fluoxetine. We demonstrate that the anti-inflammatory effect of fluoxetine coincides with intracellular lipid accumulation in human MoDMs and murine BMDMs. On the other hand, BMDMs either deficient in the LDL receptor or exposed to inflammatory OxPLs displayed an opposite pro-inflammatory response when treated with fluoxetine. Collectively, these findings show a link between the macrophage lipid landscape and fluoxetine’s immunomodulatory ability.

## METHODS

### Experimental set-up

To investigate the effects of fluoxetine on inflammatory and lipid-related responses in macrophages, a series of three related *in vitro* experiments was performed using murine BMDMs from wild-type (*Wt*) and *Ldlr /* mice, as well as normal, healthy human MoDMs. All experiments followed a similar general workflow consisting of macrophage stimulation with lipopolysaccharide (LPS), subsequent treatment with fluoxetine and/or oxidized 1- palmitoyl-2-arachidonoyl-sn-glycero-3-phosphocholine (oxPAPC, Avanti Polar Lipids 870604P), and assessment of gene expression, protein expression, cytokine secretion, and lipid accumulation. The individual experimental designs differed in stimulation sequence, treatment combinations, and cell type, and are detailed in the supplementary material. RNA was stored at −80°C, while cell culture supernatants were stored at −20°C.

### Isolation and culture of murine bone marrow-derived macrophages

BMDMs were isolated from the tibiae and femurs of wild-type (*Wt*) and *Ldlr ^-/-^* C57BL/6J mice. The final euthanasia of the experimental animals is carried out under isoflurane anesthesia with subsequent cervical dislocation. All animal studies performed were approved by the local ethical committee (approval number 30379A4). All procedures are in line with the guidelines from Directive 2010/63/EU of the European Parliament on the protection of animals used for scientific purposes. Cells were cultured in RPMI-1640 (Roswell Park Memorial Institute-1640, GIBCO Invitrogen, Breda, the Netherlands) enriched with 10% heat-inactivated fetal calf serum (iFCS, Bodinco B.V. Alkmaar, The Netherlands), penicillin (100 U/ml), streptomycin (100 μg/ml) and L-glutamine (2 mM)(all GIBCO Invitrogen, Breda, The Netherlands), supplemented with 20% L929-conditioned medium (LCM) for 8–9 days to generate BMDMs.

### Isolation and culture of human monocyte-derived macrophages

Peripheral blood mononuclear cells (PBMCs) were isolated from healthy volunteers (University Hospital RWTH Aachen, Germany) using leukocyte reduction system cones and density centrifugation as previously described [23]. In summary, PBMCs were extracted through density gradient centrifugation using Ficoll-Plaque PLUS (GE Healthcare Biosciences) and subsequently washed three times with cold phosphate-buffered saline (PBS, GIBCO Invitrogen, Breda, The Netherlands). Lastly, these isolated PBMCs were frozen in liquid nitrogen. After thawing, primary human CD14-positive monocytes were separated from PBMCs by positive selection using beads (Cat #130-097-052, Miltenyi Biotec, Leiden, The Netherlands), according to the manufacturer’s protocol. These isolated monocytes were then cultured in Dutch-modified RPMI-1640 (GIBCO Invitrogen, Breda, The Netherlands) enriched with 10% iFCS, gentamycin (5 mg/ml), pyruvate (100 mM), and Glutamax (200 mM) (all GIBCO Invitrogen, Breda, The Netherlands) supplemented with 20% LCM- conditioned medium.

### Enzyme-Linked ImmunoSorbent Assay (ELISA)

An ELISA kit was used to assess the protein expression of IL1β. Specifically, the ELISA for mouse IL1β (ab100705; Abcam) and human IL1β (ab21402; Abcam) were performed on the culture medium of BMDMs and MoDMs, respectively, according to the manufacturer’s protocol.

### RNA isolation, cDNA synthesis and quantitative real-time polymerase chain reaction

Total RNA was isolated from cultured murine BMDMs and human MoDMs with TriReagent (T9424, Sigma-Aldrich, Zwijndrecht, The Netherlands) according to the manufacturer’s instructions. The concentration and purity of the extracted RNA were assessed using a NanoDrop 2000c spectrophotometer (Thermo Fisher Scientific). According to the manufacturer’s protocol, this RNA was converted into first-strand complementary DNA (cDNA) with the SensiFASTTM cDNA synthesis kit (BIO-65054, Meridian Memphis, Tennessee). Gene expression was determined by quantitative RT-PCR (qRT-PCR) on the LightCycler® 480 (Roche, The Netherlands) using 2 ng/µl cDNA and the SensiMix SYBR & Fluorescein kit (QT615-05, Meridian, Memphis, Tennessee). A non-template and cDNA control were included for every gene. Data from qRT-PCR were analyzed using the LinRegPCR software (Version 2015.3) and normalized to the expression of housekeeping genes syntaxin 5A (Stx5a), heterogeneous nuclear ribonucleoprotein A/B (Hnrnpab), cyclophilin (Cyclo), Glyceraldehyde-3-phosphate dehydrogenase (Gapdh), and Ribosomal protein S12 (S12). Primer sequences are described in Supplementary Table 1 and 2.

### Oil Red O (ORO) staining

Murine BMDMs and human MoDMs were cultured on glass coverslips and fixed with 4% paraformaldehyde (PFA, P6148-500G, Sigma-Aldrich, Zwijndrecht, The Netherlands). Cells were subsequently stained with 1.5% ORO in isopropanol (I9515, Sigma-Aldrich, Zwijndrecht, The Netherlands) for 10 min. Representative images were acquired using a Nikon Eclipse E800 microscope (Nikon Instruments Inc.). For quantitative analysis, ORO was eluted using isopropanol and absorbance was subsequently measured at 510 nm using a Benchmark microplate reader (BioRad, Hercules, CA, USA). To account for differences in cell number between conditions, ORO absorbance was normalized to total cellular protein content as determined by Sulforhodamine B staining (not published).

### Data analysis

Statistical analysis was performed using the GraphPad Prism 8 software (GraphPad Software Inc, San Diego, CA, U.S.). The Shapiro-Wilk test, the Kolmogorov-Smirnov test, and Q-Q plots (not published) were performed to assess normality. As all data followed a normal distribution, only parametric tests were executed. Data were analyzed using a two-way ANOVA with Tukey multiple comparisons to compare between two or more groups, respectively. All quantitative data were presented as mean ± standard error of the mean (SEM). All data are the result of at least 5 independent experiments. *p<0.05 was considered statistically significant. Outlier detection was performed using the ROUT method (Q=1%) in Graphpad Prism 8 software and excluded from further analysis.

## RESULTS

### Fluoxetine dampens inflammation in wild-type macrophages while inducing lipid accumulation

To determine the immunomodulatory potential of fluoxetine, we first examined, under baseline conditions, how fluoxetine affects the immunological response of LPS-primed BMDMs and human MoDMs (Fig. 1A). Fluoxetine treatment significantly reduced IL1β cytokine production in LPS-primed murine BMDMs and human MoDMs, but not in control-primed conditions (Fig 1B-C). Similarly, gene expression levels of the pro-inflammatory markers *Il1b* and *Cxcl1* decreased significantly in LPS-primed murine BMDMs that were treated with fluoxetine, while no effect was observed for *Tnf*α and *Cxcl2* (Fig. 1D). These inflammatory findings were confirmed in LPS-primed human MoDMs, demonstrating that fluoxetine decreased the expression levels of *IL1B*, *TNF*α, and *CXCL1,* while the decrease of *CXCL2* did not reach significance (Fig. 1E). Overall, these observations show the ability of fluoxetine to dampen the LPS-induced inflammatory response.

**Figure 1:**
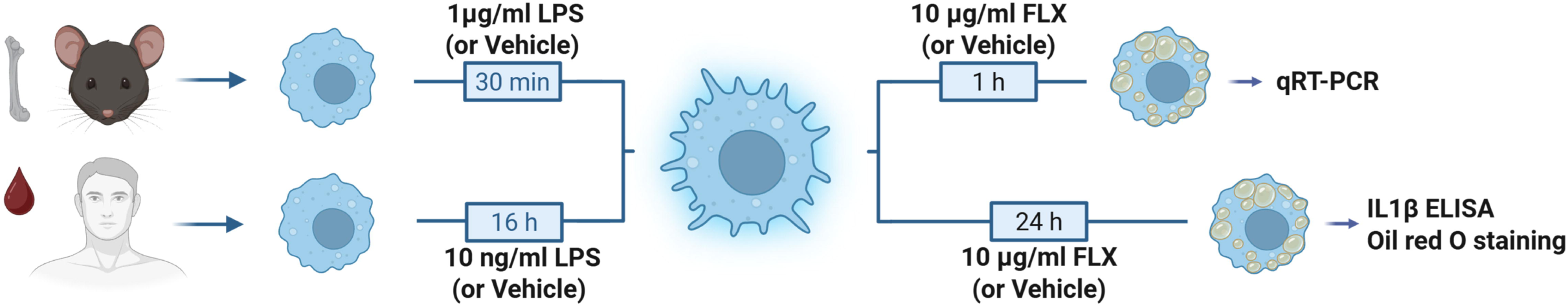
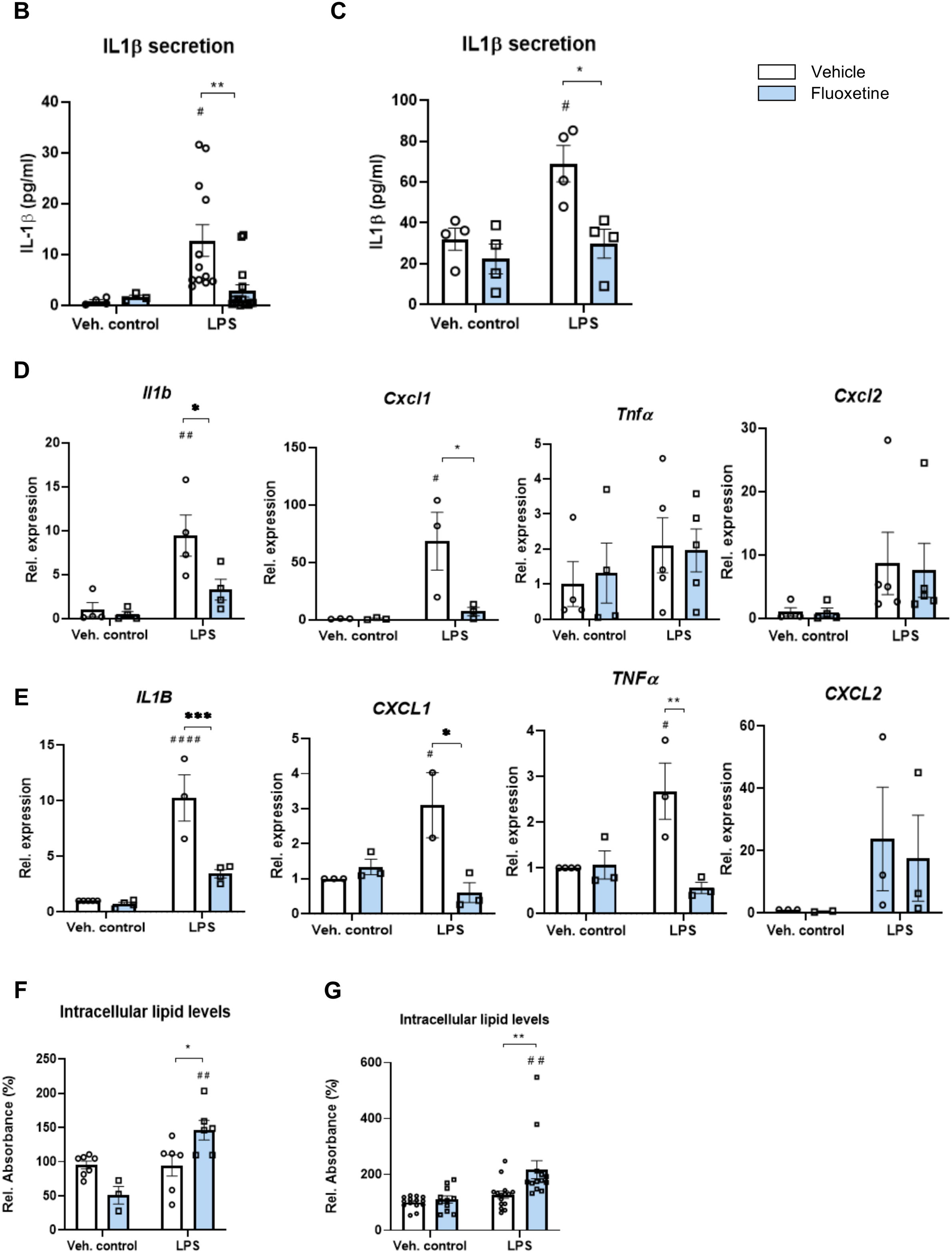
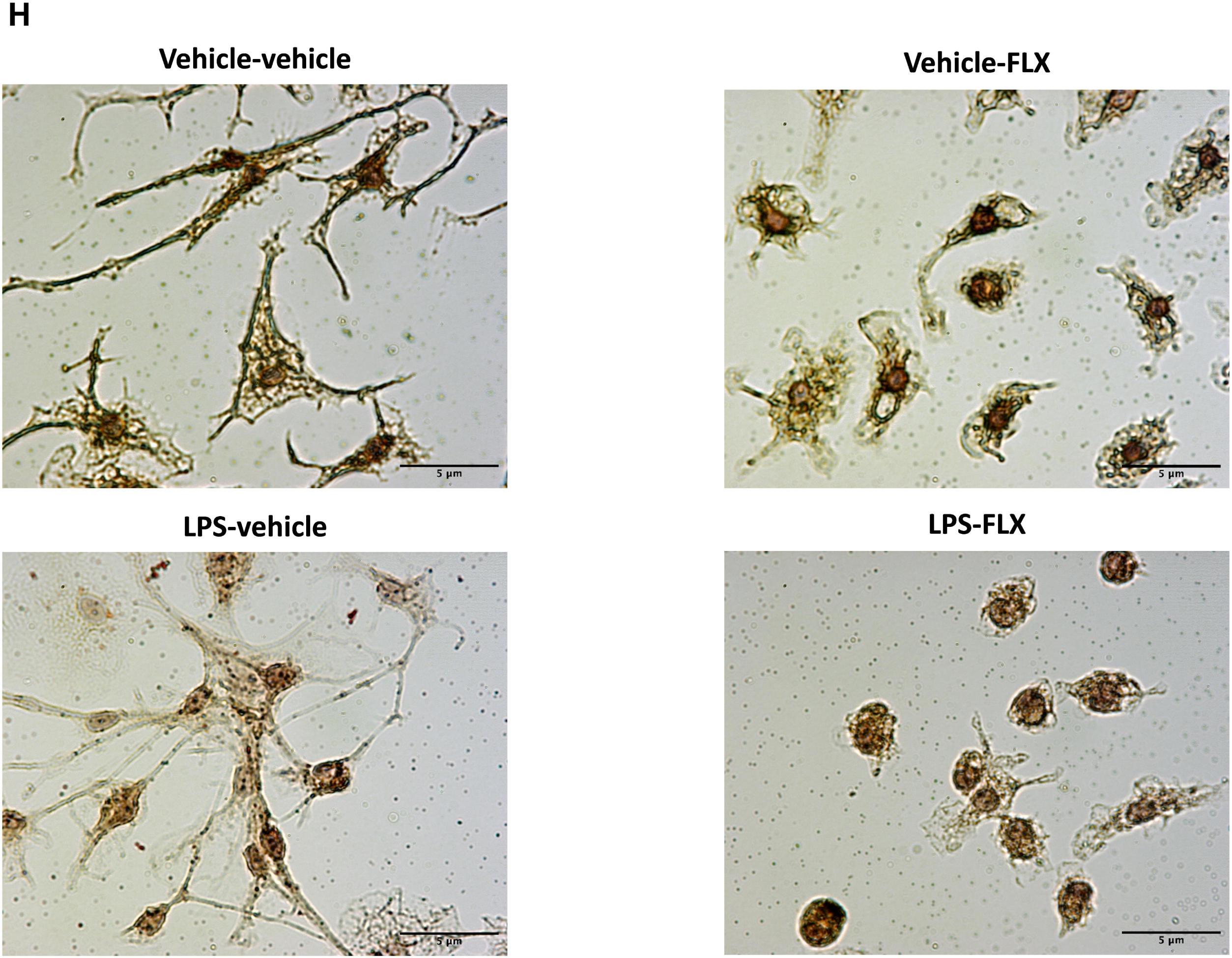
The pharmacological antidepressant fluoxetine modulates inflammation and lipid accumulation in *Wt* macrophages. A) Schematic overview of the experimental design (Created by Biorender.com). Murine *Wt* BMDMs (B and D) or MoDMs (C and E) from healthy volunteers were primed with LPS (1 µg/ml or 10 ng/ml resp.) or vehicle control for 30 minutes or 16hr resp., followed by treatment with or without fluoxetine (FLX; 10 µg/ml). B, C) IL1β protein levels in BMDMs (B) and MoDMs (C). D, E) Gene expression levels of inflammatory markers in BMDMs (D) and MoDMs (E). F, G) Quantification of Oil red O staining of fluoxetine-treated BMDMs (F) and MoDMs (G). H, I) Representative images of Oil red O staining of BMDMs (H). *p≤0.05, ** p≤0.01 and *** p≤0.001 compared with vehicle control-treated BMDMs or MoDMs by use of two-way ANOVA with Tukey’s *post hoc* correction. ^#^ p≤0.05, ^#^ ^#^ p≤0.01 and ^#^ ^#^ ^#^ ^#^ p≤0.0001 compared with vehicle control-primed BMDMs or MoDMs by use of two-way ANOVA with Tukey’s *post hoc* correction. All data were obtained from five to thirteen independent experiments. Bar graphs represent mean ± SEM. BMDM: bone marrow-derived macrophages; MoDMs: human monocyte-derived macrophages; LPS: lipopolysaccharide; IL1β: interleukin-1 beta

As lipid metabolism is meticulously regulated in inflammatory macrophages to control foam cell formation [25, 26], we performed a foam cell formation assay in macrophages via an Oil red O staining, identifying intracellular lipid levels. While fluoxetine treatment did not influence intracellular lipid levels in vehicle control-primed macrophages, intracellular lipid levels significantly increased in fluoxetine-treated, LPS-primed BMDMs (Fig. 1F + 1H). Moreover, an identical observation was made in MoDMs: fluoxetine treatment led to pronounced intracellular lipid accumulation in LPS-primed MoDMs (Fig. 1G). These findings therefore show that fluoxetine treatment of pro-inflammatory macrophages leads to reduced inflammatory responses, alongside pronounced intracellular lipid accumulation.

### Loss of the low-density lipoprotein receptor rewires the inflammatory response of pro-inflammatory BMDMs to fluoxetine

Given that LDL receptor (LDLR)-mediated lipid uptake is central to macrophage foam cell formation, we next examined whether genetic deletion of LDLR alters the inflammatory outcome of fluoxetine treatment. In contrast to *Wt* BMDMs, fluoxetine treatment in *Ldlr*^*/*^ BMDMs resulted in a significant increase in IL-1β secretion (Fig. 2B) and expression levels of the respective pro-inflammatory markers *Il1b*, *Tnf*α, *Cxcl1*, and *Cxcl2* (Fig. 2C) compared to vehicle control. Consistent with this pro-inflammatory shift, *Hmox1* expression showed a declining trend in *Ldlr /* BMDMs following fluoxetine treatment (Fig. 2D), further supporting an altered inflammatory response to fluoxetine.

**Figure 2:**
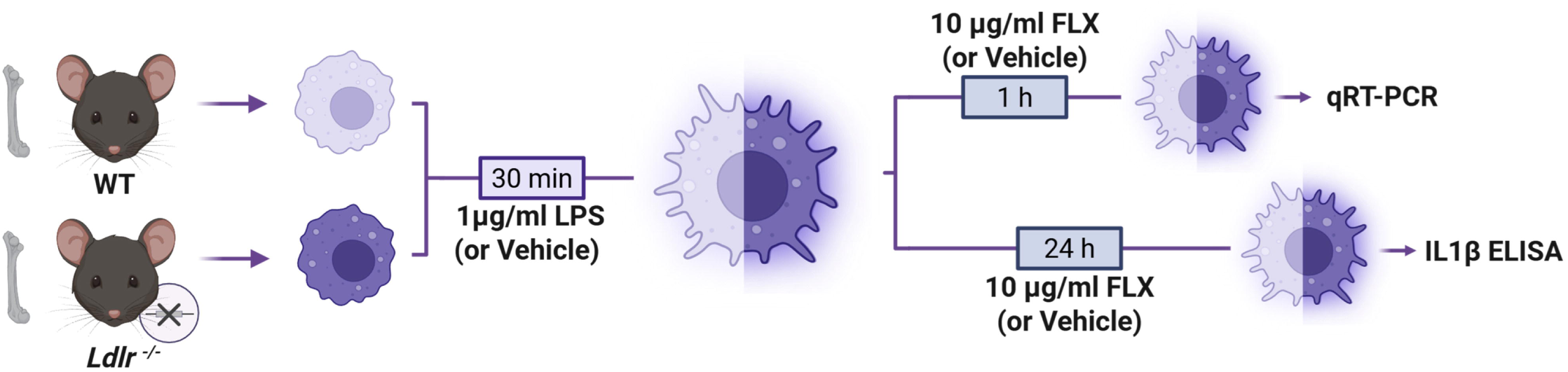
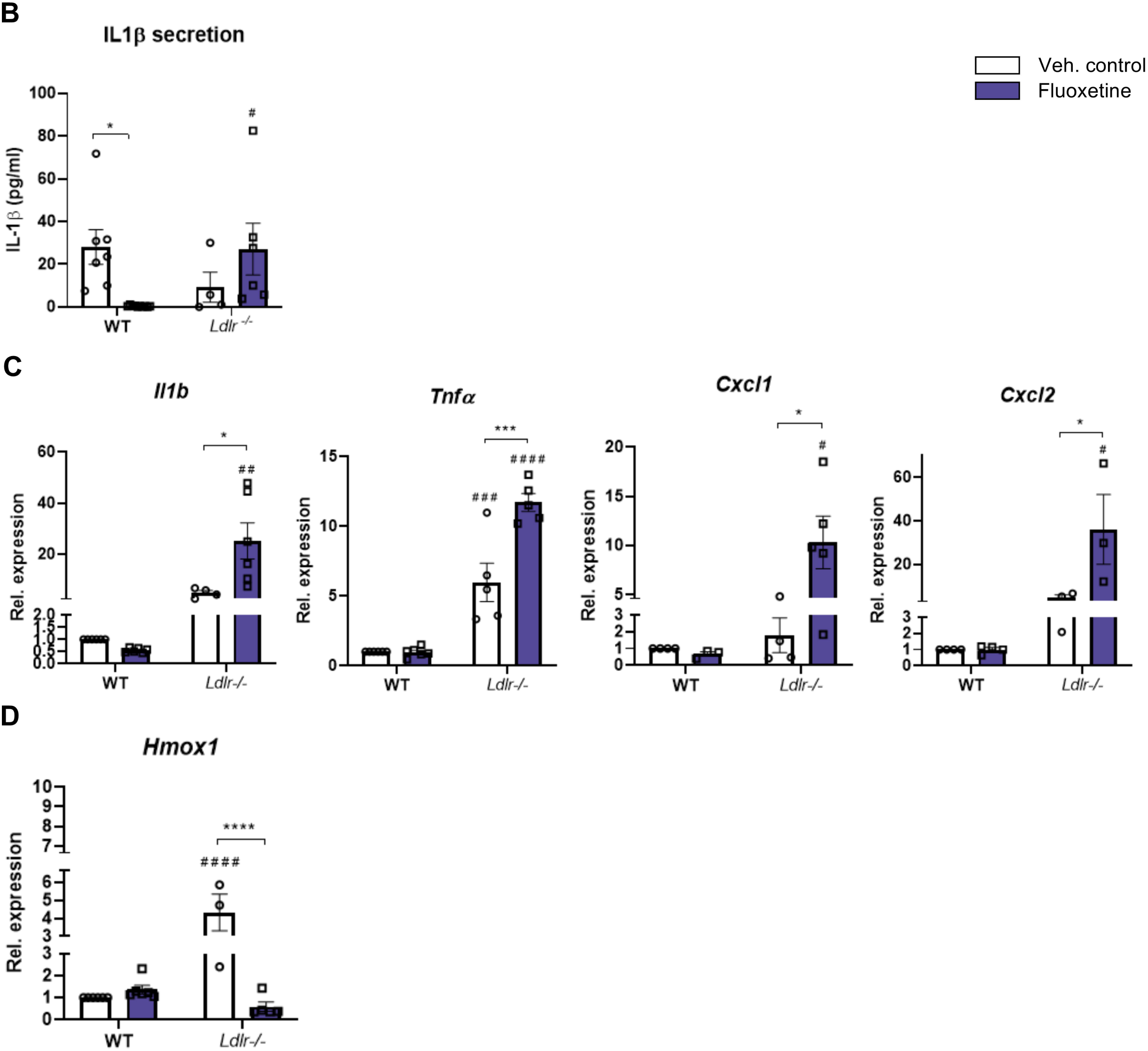
Loss of LDLR alters the inflammatory response to fluoxetine in pro- inflammatory BMDMs. A) Schematic overview of the experimental design (Created by Biorender.com). Murine *Wt* and *Ldlr^-/-^* BMDMs were primed with LPS (1 µg/ml) for 30 minutes, followed by treatment with fluoxetine (FLX; 10 µg/ml) for 1hr (gene expression analysis) or 24hr (protein expression analysis). B) IL1β protein levels in fluoxetine-treated *Wt* and *Ldlr^-/-^* BMDMs. C, D) Gene expression levels of inflammatory markers in fluoxetine- treated *Wt* and *Ldlr^-/-^*BMDMs. *p≤0.05 and *** p≤0.001 compared with vehicle control by use of two-way ANOVA with Tukey’s *post hoc* correction. ^#^ p≤0.05, ^#^ ^#^ p≤0.01, ^#^ ^#^ ^#^ p≤0.001 and ^#^ ^#^ ^#^ ^#^ p≤0.0001 compared with *Wt* BMDMs by use of two-way ANOVA with Tukey’s *post hoc* correction. Bar graphs represent mean ± SEM. BMDM: bone marrow-derived macrophages; LPS: lipopolysaccharide; IL1β: interleukin-1 beta; Tnfα: tumor necrosis factor alpha; Cxcl1: chemokine (C-X-C-motif)ligand 1; Cxcl2: chemokine (C-X-C-motif)ligand 2; Hmox1: heme oxygenase 1

Moreover, two-way ANOVA analysis revealed a significant LDLR status × fluoxetine treatment interaction across all analyzed inflammatory markers (Table 1), indicating that absence of LDLR significantly alters the macrophage inflammatory response to fluoxetine. Together, these data demonstrate that LDLR status critically determines the direction of the inflammatory response to fluoxetine in pro-inflammatory BMDMs.

**Table 1:**
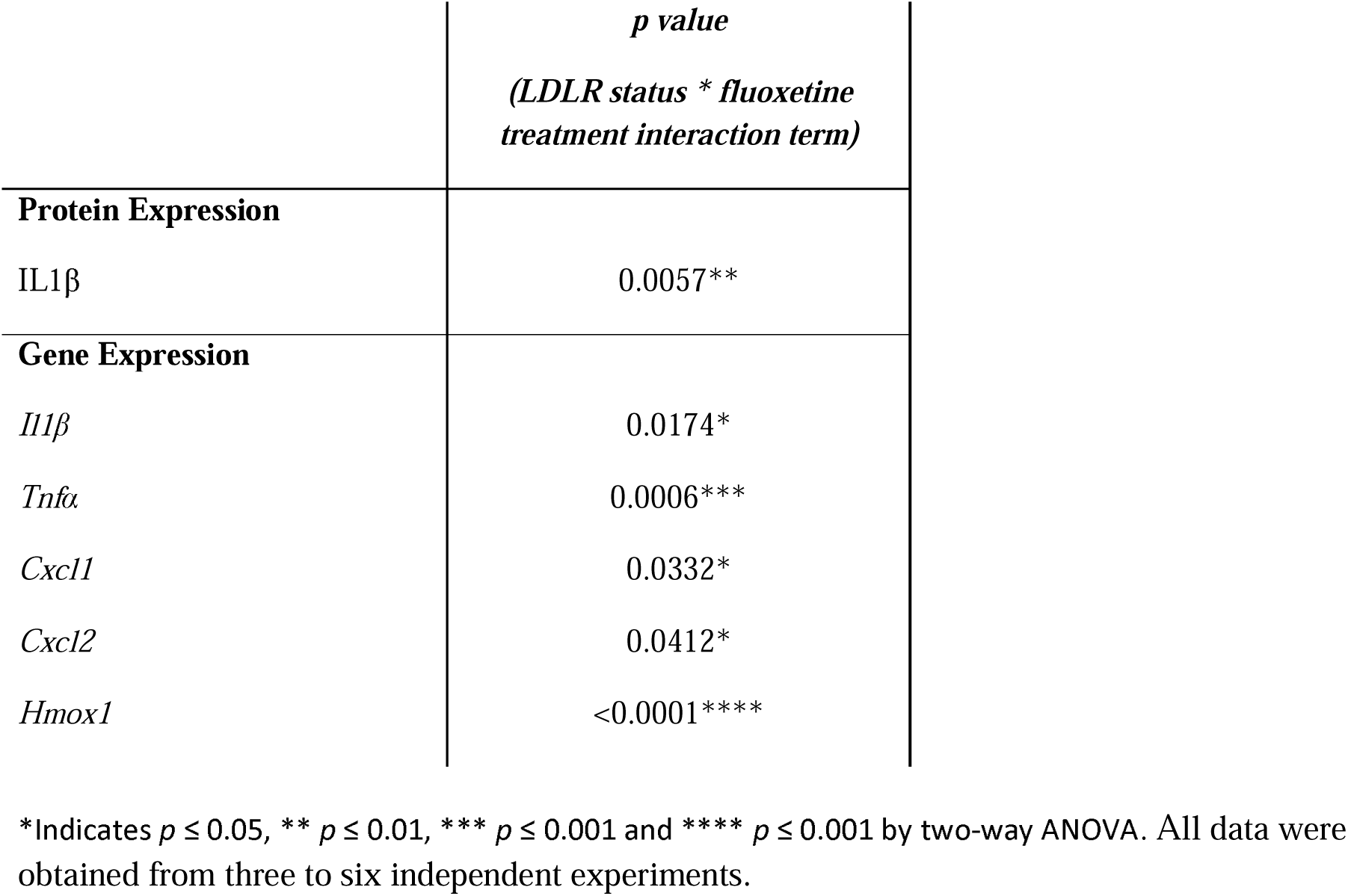
LDLR status × fluoxetine treatment interactions demonstrate regulation of fluoxetine- induced inflammatory responses by LDLR status.

### An OxPAPC-rich environment reverses the inflammatory response of pro-inflammatory BMDMs to fluoxetine

To determine whether the reversal of fluoxetine’s inflammatory effects in BMDMs also occurs in lipid-rich inflammatory environments, such as those associated with obesity [27], we next examined BMDM responses to fluoxetine in the presence of inflammatory oxidized phospholipids. In line with our hypothesis, exposing *Wt* BMDMs to an OxPAPC-rich environment similarly resulted in an opposite, pro-inflammatory effect of fluoxetine, as evidenced by increased IL-1β protein levels (Fig. 3B) and gene expression levels of the inflammatory markers *Il1b*, *Cxcl1*, *Cxcl2 and Hmox1* (Fig. 3C and 3D). Moreover, while *Tnf*α changes did not reach significance, two-way ANOVA analysis revealed a significant OxPAPC × fluoxetine treatment interaction across nearly all analyzed inflammatory markers (Table 2), supporting the claim that an OxPAPC-rich environment alters the macrophage inflammatory response to fluoxetine.

**Figure 3:**
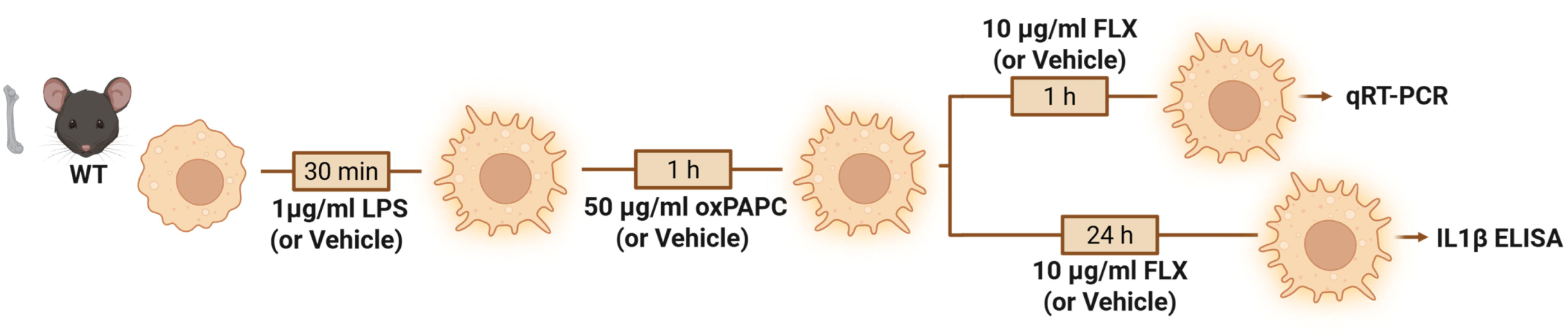
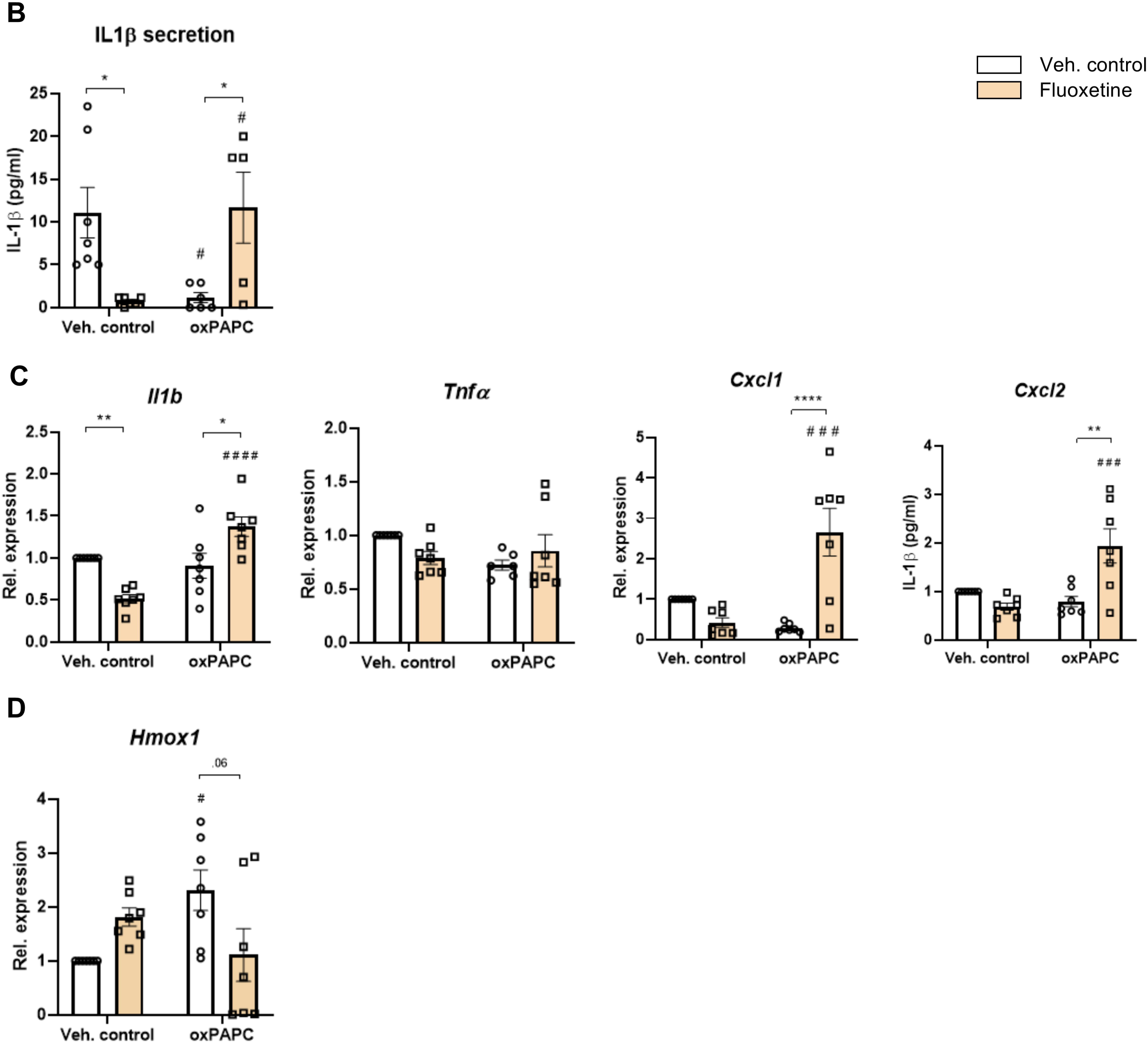
An oxPAPC-rich environment influences the inflammatory response to fluoxetine in pro-inflammatory BMDMs. A) Schematic overview of the experimental design (Created by Biorender.com). Murine *Wt* BMDMs were primed with LPS (1 µg/ml) for 30 minutes, followed by exposure to oxPAPC (50 µg/ml) for 1hr to establish an oxPAPC-rich environment. Subsequently, BMDMs were treated with fluoxetine (FLX; 10 µg/ml) for 1hr (gene expression analysis) or 24hr (protein expression analysis). For more detailed information regarding the experimental design, we refer to the Material and Methods. B) IL1β protein levels in fluoxetine-treated *Wt* BMDMs in an oxPAPC-rich environment. C-D) Gene expression levels of inflammatory markers in fluoxetine-treated *Wt* BMDMs in an oxPAPC- rich environment. *p≤0.05, ** p≤0.01 and **** p≤0.01 compared with vehicle control by use of two-way ANOVA with Tukey’s *post hoc* correction. ^#^ p≤0.05, ^#^ ^#^ ^#^ p≤0.001 and ^#^ ^#^ ^#^ ^#^ p≤0.0001 compared vehicle control by use of two-way ANOVA with Tukey’s *post hoc* correction. Bar graphs represent mean ± SEM. BMDM: bone marrow-derived macrophages; LPS: lipopolysaccharide; IL1β: interleukin-1 beta; Tnfα: tumor necrosis factor alpha; Cxcl1: chemokine (C-X-C-motif)ligand 1; Cxcl2: chemokine (C-X-C-motif)ligand 2; Hmox1: heme oxygenase 1; oxPAPC: Oxidized 1-palmitoyl-2-arachidonyl-sn-glycero-3-phosphorylcholine.

**Table 2:** OxPAPC presence × fluoxetine treatment interactions demonstrate regulation of fluoxetine-induced inflammatory responses by an oxPAPC-rich environment.

|  | *p value*<br>(*OxPAPC presence \* fluoxetine treatment interaction term*) |
| --- | --- |
| **Protein Expression** |  |
| IL1β | 0.0004*** |
| **Gene Expression** |  |
| *Il1β* | <0.0001**** |
| *Tnfa* | 0.0587 |
| *Cxcl1* | <0.0001**** |
| *Cxcl2* | 0.0007*** |
| *Hmox1* | 0.0113* |

\*Indicates $p \leq 0.05$ , \*\*\* $p \leq 0.001$ and \*\*\*\* $p \leq 0.001$ by two-way ANOVA. All data were obtained from three to seven independent experiments.

## DISCUSSION

As the primary orchestrators of innate immunity, macrophages possess extraordinary functional plasticity that is increasingly recognized as a factor in the systemic inflammation associated with depressive disorders [28, 29]. While the ability of SSRIs such as fluoxetine to dampen this inflammatory response is well-documented [30], the metabolic requirements that shape this therapeutic effect remain largely unexplored. Here, we demonstrate that the macrophage lipid landscape and the inflammatory response to fluoxetine are dynamically linked, suggesting that the drug’s immunomodulatory efficacy is not an isolated signaling event. Instead, our findings suggest that the lipid environment and subsequent cellular processing of lipids act as a critical ‘rheostat,’ where the metabolic state of the macrophage fundamentally determines fluoxetine’s anti-inflammatory impact.

Our results demonstrate that fluoxetine treatment reduces inflammation while simultaneously promoting intracellular lipid accumulation, as evidenced by increased Oil Red O staining in both murine and human macrophages. This observation is particularly striking as it stands in direct contrast to the traditional paradigm, in which lipid-laden macrophages, typically associated with a ‘foamy’ phenotype, are viewed as inherently pro-inflammatory and detrimental to tissue homeostasis [9, 31, 32]. Therefore, the ability of fluoxetine to suppress inflammation while promoting intracellular lipid levels raises the question of whether this SSRI reconfigures macrophages to maintain such a quiescent inflammatory state. A potential answer to this question may lie in fluoxetine’s ability to modulate lysosomal function and lipid processing. Specifically, fluoxetine belongs to the class of cationic amphiphilic drugs (CADs), which are characterized by their significant accumulation within subcellular organelles, particularly the lysosomal compartment [33]. This localization is relevant to the inflammatory response as lysosomal membrane permeabilization (LMP) is a critical trigger for inflammasome activation [34]. Previous studies have shown that the accumulation of unesterified cholesterol can stabilize the lysosomal membrane, thereby protecting against LMP and reducing the release of pro-inflammatory cytokines [35, 36]. As such, given that CADs have been implicated in promoting the lysosomal sequestration of unesterified cholesterol, in our study, fluoxetine may suppress inflammation by reinforcing lysosomal integrity and preventing the release of pro-inflammatory signals. However, it is important to note that our intracellular lipid measurements specifically quantified neutral lipids, such as cholesterol esters and triglycerides. This finding therefore suggests a broader metabolic shift upon fluoxetine treatment than the sequestration of free unesterified cholesterol alone. This is in line with reports describing increased plasma and hepatic triglyceride levels in mice treated with fluoxetine [37] as well as depressive patients developing moderate [37] to severe [38, 39] hypertriglyceridemia upon fluoxetine treatment. Ultimately, these clinical parallels reinforce the notion that the anti-inflammatory effect of fluoxetine is associated with a systemic and cellular reconfiguration of lipids.

The observation that fluoxetine’s anti-inflammatory effect is reversed in LDLR-deficient macrophages suggests that an intact LDL receptor is required for fluoxetine’s anti-inflammatory action. The LDL receptor facilitates the delivery of LDL-derived cholesterol to the lysosomal system. Our findings therefore reinforce the hypothesis that the lysosomal sequestration of unesterified cholesterol is an underlying mechanism explaining fluoxetine’s anti-inflammatory effects [35]. Specifically, in the absence of LDLR-driven lipid supply, the macrophage is deprived of the lipid substrates required for fluoxetine to stabilize the lysosomal membrane. This decoupling hampers the lysosomal integrity, thereby failing to suppress the subsequent inflammatory signaling. Moreover, clinically, a deficiency in the LDL receptor is associated with patients diagnosed with familial hypercholesterolemia (FH) [14], a disorder characterized by elevated plasma LDL cholesterol and total cholesterol levels. While the rate of fluoxetine nonresponse in FH patients is (to our knowledge) currently unknown, the association between elevated plasma cholesterol levels and poor fluoxetine response in MDD [12] appears to confirm a role for cholesterol metabolism in fluoxetine’s efficacy. By suggesting LDL receptor dysfunction as a contributing factor to fluoxetine nonresponse, this study provides a rationale for expanding research into the role of the LDL receptor in antidepressant sensitivity/resistance. Investigating these cholesterol-dependent pathways may also offer deeper insight into the underlying biological mechanisms of depressive disorders.

Also, we uncovered that fluoxetine’s anti-inflammatory effect can be subverted by the nature of the local lipid environment. Specifically, the presence of OxPLs, a highly reactive and pro-inflammatory lipid species [20], reversed the anti-inflammatory properties of fluoxetine. This observation highlights a potential metabolic boundary for fluoxetine’s efficacy in metabolic syndrome-related conditions such as MASH [21, 32] and atherosclerosis [22, 40], where OxPLs are known to accumulate. Our findings suggest that in these lipotoxic environments, the pro-inflammatory signaling of OxPLs have the ability to neutralize fluoxetine’s protective effects. This hypothesis is confirmed by studies showing that depressed patients with an unfavorable metabolic phenotype (i.e. increased BMI, dyslipidemia, and increased systemic inflammatory parameters) demonstrate worse response rates towards antidepressant monotherapies [41–43]. Therefore, addressing lipid (immuno-)metabolic dysfunction (with OxPLs as a potential direct target) as a core determinant of antidepressant treatment resistance in future research could therefore be instrumental in identifying the metabolic profiles most favorable to antidepressant success.

To our knowledge, we are the first to show a coupling between lipid metabolism and the inflammatory response of fluoxetine at the macrophage level. However, several limitations in our study must be acknowledged. While our findings are validated in both murine and human primary macrophages *in vitro*, our study lacks corroborating evidence from *in vivo* models or clinical samples. Specifically, analyzing macrophage phenotypes from depressed individuals, comparing those treated with fluoxetine to untreated controls, remains a necessary next step to confirm the physiological relevance of the lipid-inflammatory axis. Furthermore, while the link between systemic inflammation and depression is well-established, our current data (and experimental design) do not directly address whether the cellular metabolic shifts correlate with improved behavioral or clinical outcomes in depression.

In summary, our findings suggest that the anti-inflammatory response of fluoxetine is linked to lipid metabolism in macrophages. The extent to which this metabolic interplay dictates treatment outcomes in depressed individuals remains to be established in future studies.

## ACKNOWLEDGEMENTS

Certain figures were created with BioRender.com. LLMs (ChatGPT and Gemini) were used solely for language refinement and stylistic suggestions. No scientific content, data analysis, or interpretation was generated by the LLM. All final text was written, reviewed, and approved by the authors. Furthermore, the authors also like to thank Dr. Dennis Meesters for his assistance in isolating human monocyte-derived macrophages.

## AUTHOR CONTRIBUTIONS

AJ, JG, LT, EALB, EvdV and TH were responsible for conceptualization of the experiments. AJ, JG, DM, TH performed the experiments. Data analysis was performed by AJ, JG and JG, RSS and TH wrote the draft manuscript, which was reviewed and edited by all co-authors.

