## Supplementary Table 1 for "The lipid landscape shapes the immunomodulatory potential of fluoxetine in macrophages"

**Supplementary Table 1:** Overview of murine primers used for quantitative RT-PCR.

| **Gene** | **Primer type** | **Primer sequence** |
| --- | --- | --- |
| *Cyclo* | Forward Reverse | 5’ TTCCTCCTTTCACAGAATTATTCCA 3’  5’ CCGCCAGTGCCATTATGG 3’ |
| *S12* | Forward Reverse | 5’ GGAAGGCATAGCTGCTGGAGGTGT 3’  5’ CCTTCGATGACATCCTTGGCCTGAG 3’ |
| *Stx5a* | Forward Reverse | 5' CTGAAACAGCAGAGGAACCGTC 3'  5' GGTCCATCATGTCAATAGCCACG 3' |
| *Hnrnpab* | Forward Reverse | 5' CCAACACTGGACGATCAAGAGG 3'  5' ATGACACGACCATCCAGCCTGT 3' |
| *Cxcl1* | Forward Reverse | 5’ TCCAGAGCTTGAAGGTGTTGCC 3’  5’ AACCAAGGGAGCTTCAGGGTCA 3’ |
| *Cxcl2* | Forward Reverse | 5’ CGCTGTCAATGCCTGAAGAC 3’  5’ ACACTCAAGCTCTGGATGTTCTTG 3’ |
| *Tnfα* | Forward Reverse | 5’ CATCTTCTCAAAATTCGAGTGACAA 3’  5’ TGGGAGTAGACAAGGTACAACCC 3’ |
| *Il1b* | Forward Reverse | 5’AAAGAATCTATACCTGTCCTGTGTAATGAAA 3’  5’ GGTATTGCTTGGGATCCACACT 3’ |
| *Hmox1* | Forward Reverse | 5’ CCGCCTTCCTGCTCAACAT 3’  5’ ATCTGTGAGGGACTCTGGTCTTTG 3’ |

*Cyclo, Cyclophilin; S12, Ribosomal protein S12; Stx5a, Syntaxin 5A; Hnrnpab; Heterogeneous nuclear ribonucleoprotein A/B; Cxcl1/2, C-X-C motif ligand 1/2; Tnfa; Tumor necrosis factor* α*; Il1b, Interleukin 1ß; Hmox1, Heme oxygenase 1*
