## Supplementary Table 2 for "The lipid landscape shapes the immunomodulatory potential of fluoxetine in macrophages"

**Supplementary Table 2:** Overview of human primers used for quantitative RT-PCR.

| **Gene** | **Primer type** | **Primer sequence** |
| --- | --- | --- |
| *GAPDH* | Forward Reverse | 5’ TTCAACAGCGACACCCACT 3’  5' TTCCTCTTGTGCTCTTGCT 3' |
| *STX5A* | Forward Reverse | 5' GAACACGGATCAGGGTGTCTA 3'  5' ACGTTCTCGTCGATCCTCTG 3' |
| *TNFα* | Forward Reverse | 5' CTCTTCTGCCTGCTGCACTTTG 3'  5' ATGGGCTACAGGCTTGTCACTC 3' |
| *IL1B* | Forward Reverse | 5' CTGAGCTCGCCAGTGAAATG 3'  5' TTTAGGGCCATCAGCTTCAAA 3' |
| *CXCL1* | Forward Reverse | 5’ GGCGCTGTCATCGATTTCTT 3’  5’ TGGAGCTTATTAAAGGCATTCTTCA 3’ |
| *CXCL2* | Forward Reverse | 5’ GGCCTTCGATTCTGGATTCA 3’  5’ CAATTTTGTCCACGTGTTGAGATC 3’ |

*GAPDH, Glyceraldehyde-3-phosphate dehydrogenase STX5A, Syntaxin 5A; CXCLl1/2, C-X-C motif ligand 1/2; TNFα; Tumor necrosis factor* α*; IL1B, Interleukin 1ß*
