## Supplementary Material for "The lipid landscape shapes the immunomodulatory potential of fluoxetine in macrophages"

**METHODS**

**Experiment 1: Effect of FLX in WT murine BMDMs and healthy human MoDMs**

To investigate the effect of FLX under inflammatory conditions in murine and human macrophages, WT BMDMs and healthy human MoDMs were first primed with 20% LCM medium (vehicle control) or lipopolysaccharide (LPS; 1 µg/ml for 30 min in murine cells, 10 ng/ml for 16hr in human cells; L2880, Sigma-Aldrich, Zwijndrecht, The Netherlands) to mimic a pro-inflammatory environment as observed during depression [9]. Subsequently, cells were treated with FLX (10 µg/ml; F132-10MG, Sigma-Aldrich, Zwijndrecht, The Netherlands) or vehicle control for 1 hour (for gene expression analysis) or 24 hours (for protein expression and staining analyses).

**Experiment 2: Effect of FLX in pro-inflammatory *Ldlr⁻/⁻* BMDMs**

To determine whether the inflammatory effects of FLX were preserved upon loss of the low-density lipoprotein receptor (LDLR), BMDMs derived from *Ldlr⁻/⁻* mice were primed with LPS (1 µg/ml) for 30 min. After priming, cells were washed with PBS and treated with either FLX (10 µg/ml) or 20% LCM medium (vehicle control) for 1 hour (for gene expression analysis) or 24 hours (for protein expression and staining analyses).

**Experiment 3: Effect of FLX in OxPAPC-exposed pro-inflammatory WT BMDMs**
To investigate the inflammatory effect of FLX on macrophage responses to oxidized phospholipids, WT BMDMs were first primed with lipopolysaccharide (LPS; 1 µg/ml; L2880, Sigma-Aldrich, Zwijndrecht, The Netherlands) or 20% LCM medium (vehicle control) for 30 minutes. Following priming, cells were treated with oxidized 1-palmitoyl-2-arachidonyl-sn-glycero-3-phosphorylcholine (OxPAPC; 50 µg/ml; Avanti Polar Lipids 870604P, Sigma-Aldrich, Zwijndrecht, The Netherlands) or 20% LCM medium (vehicle control) for 1 hour. Subsequently, cells were exposed to OxPAPC (50 µg/ml) in combination with fluoxetine (FLX; 10 µg/ml; F132-10MG, Sigma-Aldrich, Zwijndrecht, The Netherlands), OxPAPC alone, FLX alone, or 20% LCM medium (vehicle control) for either 1 hour (for gene expression analysis) or 24 hours (for protein expression analysis).
